# Persistent tailoring of MSC activation through genetic priming

**DOI:** 10.1101/2024.02.01.578489

**Authors:** Michael A. Beauregard, Guy C. Bedford, Daniel A. Brenner, Leonardo D. Sanchez Solis, Tomoki Nishiguchi, Abhimanyu, Santiago Carrero Longlax, Barun Mahata, Omid Veiseh, Pamela L. Wenzel, Andrew R. DiNardo, Isaac B. Hilton, Michael R. Diehl

## Abstract

Mesenchymal stem/stromal cells (MSCs) are an attractive platform for cell therapy due to their safety profile and unique ability to secrete broad arrays of immunomodulatory and regenerative molecules. Yet, MSCs are well known to require preconditioning or priming to boost their therapeutic efficacy. Current priming methods offer limited control over MSC activation, yield transient effects, and often induce expression of pro-inflammatory effectors that can potentiate immunogenicity. Here, we describe a ‘genetic priming’ method that can both selectively and sustainably boost MSC potency via the controlled expression of the inflammatory-stimulus-responsive transcription factor IRF1 (interferon response factor 1). MSCs engineered to hyper-express IRF1 recapitulate many core responses that are accessed by biochemical priming using the proinflammatory cytokine interferon-γ (IFNγ). This includes the upregulation of anti-inflammatory effector molecules and the potentiation of MSC capacities to suppress T cell activation. However, we show that IRF1-mediated genetic priming is much more persistent than biochemical priming and can circumvent IFNγ-dependent expression of immunogenic MHC class II molecules. Together, the ability to sustainably activate and selectively tailor MSC priming responses creates the possibility of programming MSC activation more comprehensively for therapeutic applications.

**GRAPHICAL ABSTRACT:** 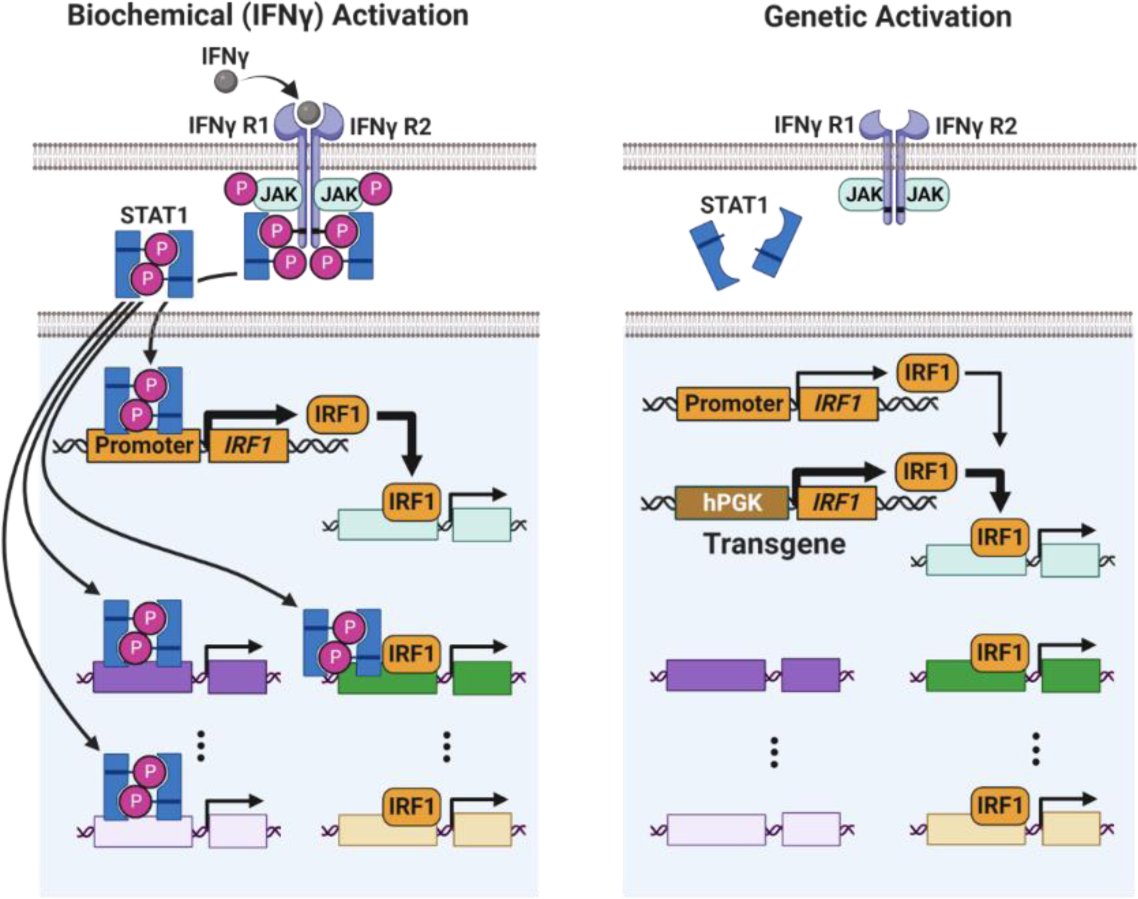

## INTRODUCTION

Mesenchymal Stem/Stromal Cells (MSCs) are widely recognized for their potential for treating diverse classes of human diseases and disorders.^1^ The efficacy of MSCs or their cell-free products^2,3^ has been studied in hundreds of clinical trials^4^ across multiple settings spanning neurodegenerative disorders,^5^ inflammatory bowel disease,^6^ cardiac diseases,^7^ COVID-19,^8^ and others. This broad potential stems from the ability of MSCs to simultaneously express and secrete a spectrum of different pleiotropic immunomodulatory and regenerative chemokines, cytokines, growth factors, metabolites, and vesicles.^9,10^ Virtually all MSC-based cell therapies seek to leverage the delivery of these effector molecules to suppress inflammation, restore immune homeostasis, and promote healing.^11,12^

Despite their promise, MSC therapies have shown limited clinical efficacy.^13^ Multiple factors contribute to this deficiency, many of which can be linked to insufficient cell potency.^4,14^ Overall, it is widely accepted that MSCs must be primed biochemically^15,16^ or biophysically^17,18^ to activate and/or enhance their production of immune effector molecules since basal unstimulated MSCs do not typically produce these molecules in sufficient quantities. Pro-inflammatory stimuli including cytokines such as IFNγ, TNFα, and IL-1β or pathogenic molecules such as lipopolysaccharide (LPS)^19^ have been explored extensively due to their ability to mimic natural priming of MSCs by activated immune cells during infections or injuries.^20^ These molecules induce comprehensive changes to the MSC transcriptome, proteome, and secretome^21^ via the activation of stimulus-responsive transcription factors (TFs).^15^ As a key example, IFNγ bolsters MSC potency through the TF STAT1 (signal transducer of activation 1).^22^ Stimulation with IFNγ activates STAT1 phosphorylation, dimerization, and translocation to the nucleus, where it binds to gamma-activated sequences (GAS) within the promoters of many different interferon-stimulated genes (ISGs).^23,24^ This includes additional TFs that can function as signaling intermediates, like IRF1 (interferon response factor 1), which further broaden transcriptional responses to IFNγ signaling by binding to interferon-stimulated response elements (ISREs) within the promoters of additional IRGs, notably with and/or without STAT1.^25^ Among other effectors, IRF1 is well known to regulate IDO1 (indoleamine 2,3-dioxygenase 1),^26,27^ which is recognized as a key determinant of the immunosuppressive potency of MSCs.^28–30^

Although biochemical and biophysical / biomechanical priming methods have both been shown to boost MSC bioactivity and improve therapeutic responses, several challenges remain that affect the functionality and durability of primed MSCs. In particular, MSC activation tends to be transient and short-lived.^31^ The effect of biochemical priming by IFNγ and other stimulants has been shown to decay within a few days in stimulus washout experiments.^32^ More persistent activation has been achieved using engineered biomaterials that integrate the slow-release of IFNγ to provide constant stimulation.^31^ Yet, biochemical stimulation is also difficult to modify and can induce the expression of unwanted and deleterious immunogenic and/or proinflammatory effectors.^33,34^ For example, in addition to other proinflammatory factors, IFNγ strongly induces the expression of the transcriptional coactivator CIITA (class II transcriptional activator).^35^ CIITA, in turn, upregulates the expression of multiple MHC class II molecules that can potentiate MSC immunogenicity and promote MSC clearance by CD4 T cells.^36^

MSCs have been genetically engineered to constitutively overexpress transgenes encoding for different effector molecules including IDO1,^37^ COX-2,^38^ and other effectors.^39^ While constitutive overexpression of these molecules can address the durability of MSC activation, current genetic engineering technologies can typically only manipulate the expression of a few genes at a time; far less than the number of genes that are activated via biochemical or biomechanical stimuli. The multifaceted secretome of MSCs, especially of primed MSCs, facilitates regulation of multiple classes of immune and immune-supporting cells,^40^ and this is among the unique properties that make MSCs attractive candidates for cell therapy.

Here, we demonstrate that stable hyperexpression of IRF1 can be leveraged to upregulate broad arrays of immunomodulatory effector genes and recapitulate the potency gains that can be achieved via traditional biochemical priming with IFNγ. However, genetic priming via IRF1 was also shown to circumvent the activation of STAT1 and key STAT1-dependent downstream ISGs like CIITA. In contrast to IFNγ-primed cells, this effect was associated with low MHC class II gene expression and improved retention of MSC hypo-immunogenicity in MSC-peripheral blood mononuclear cell (PBMC) coculture assays. Finally, sustained (>21 day) priming was also demonstrated in primary human adipose-derived MSCs (AD-MSCs), where activated IDO1 production was maintained for weeks. Together, these results demonstrate engineered expression of intermediate TFs like IRF1 can facilitate selective and persistent programming of MSCs in ways that activate large numbers of therapeutically relevant effector molecules while minimizing the activation of other molecules that have the potential to impair therapeutic efficacy. Such control may open routes to better tailor and maintain the activated phenotypes of MSCs for specific therapeutic applications.

## RESULTS

### IRF1 overexpression mimics key activation signatures of IFNγ priming

The utility of IRF1 transgenes for priming was first tested using immortalized hTERT-MSCs that were transduced (MOI = 1) with a lentiviral vector that encoded for a full length IRF1 transgene and eGFP (Figure 1A, 1B, Supplemental Figure S1-S4). The resulting IRF1-overexpressing cells (MSC^IRF1^) displayed enhanced IRF1 transcription, translation, and nuclear localization by RT-qPCR and immunocytochemistry (Figure 1C, D). Although biochemical stimulation with IFNγ for 24 hours (MSC^IFNγ^) induced higher transcription of endogenous IRF1 compared to MSC^IRF1^, both methods yielded >25,000-fold induction of IDO1 transcription (Figure 1D), which is a key ISG target of IRF1 and a signature of IFNγ priming. IDO1 exerts immunomodulatory effects by converting tryptophan into kynurenine within the tryptophan metabolic pathway.^41^ Kynurenine levels were also elevated using both priming techniques and were indistinguishable between MSC^IRF1^ and MSC^IFNγ^ (p = 0.63; Figure 1E). Together these results demonstrate that IRF1 transgene expression in MSCs can recapitulate key activation signatures of IFNγ stimulation.

**Figure 1.**
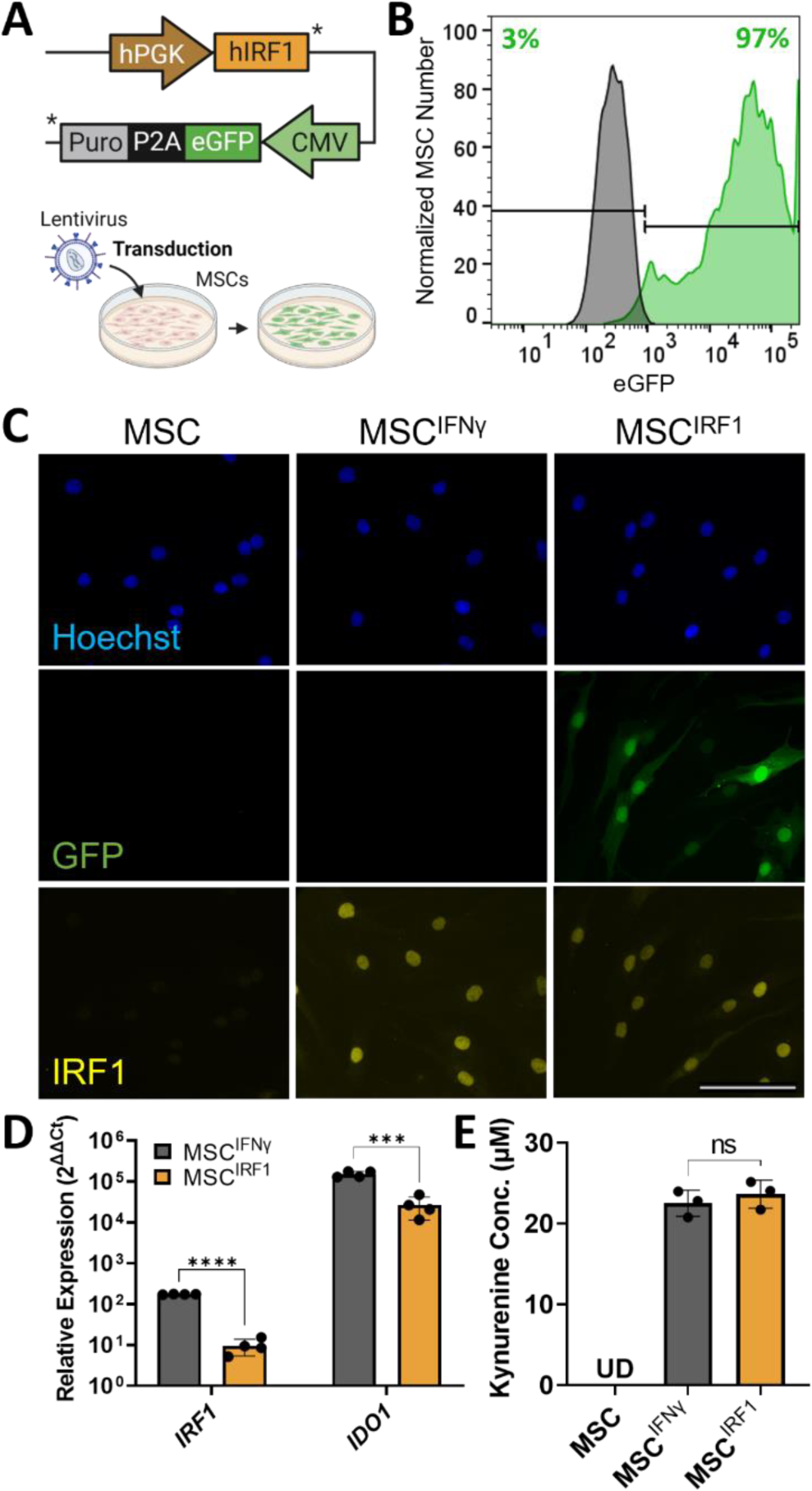
Genetic priming MSCS via overexpression of IRF1. (A) IRF1 lentiviral construct design. (B) Flow-based quantification of lentiviral transduction efficiency. (C) IRF1 transgene expression and IFNγ stimulation at 50ng/mL for 24 hours both yield nuclear localization of IRF1. IRF1 was visualized by immunocytochemical staining. Scale bar 100μm. (D and E) IRF1 overexpression and IFNγ stimulation (50 ng/mL for 24 hours) both drive IDO1 transcription (panel D) and upregulate kynurenine production (panel E). Relative expression, 2^ΔΔCt^, in RT-qPCR analyses was calculated relative to unprimed MSCs for both MSC^IFNγ^ and MSC^IRF1^. Statistical analyses were performed using multiple unpaired t tests. Kynurenine production was assayed using an Ehrlich reaction and analyzed using one way ANOVA. *p<0.05, **p < 0.01, ***p < 0.001, ****p < 0.0001 and ns, not significant. Error bars represent standard deviation.

### IRF1 activates MSC-mediated T cell suppression

We next characterized and compared IRF1 and IFNγ-induced changes to the MSC transcriptome using bulk RNA sequencing (RNAseq) (Figures 2A-2D). Both priming methods induced transcriptional changes, yielding over 1900 differentially expressed genes (DEGs), each relative to untreated control MSCs (p < 0.05; FC > 2) (Figure 2A-C). In addition to *IDO1*, upregulated DEGs included *COX-2* (*PTGS2*), a second key immunomodulatory effector enzyme that produces the anti-inflammatory small molecule prostaglandin E2^42^ (PGE2), along with multiple effector genes associated with T cell suppression including *IL4I1*,^43^ *Galactin-9*^44,45^ *(LGALS9)*, PD-L2^46^ (*PDCD1LG2*), and *FGL2*^47^ (Figure 2A). Along these lines, normalized enrichment scores from gene ontology (GO) analysis show transcription in both MSC^IRF1^ and MSC^IFNγ^ is positively enriched for genes corresponding to the GO terms ‘negative regulation of T cell proliferation’ (0042130) and ‘negative regulation of T cell activation’ (005086800) (Figure 2D, Supplemental Figure S5).

**Figure 2.**
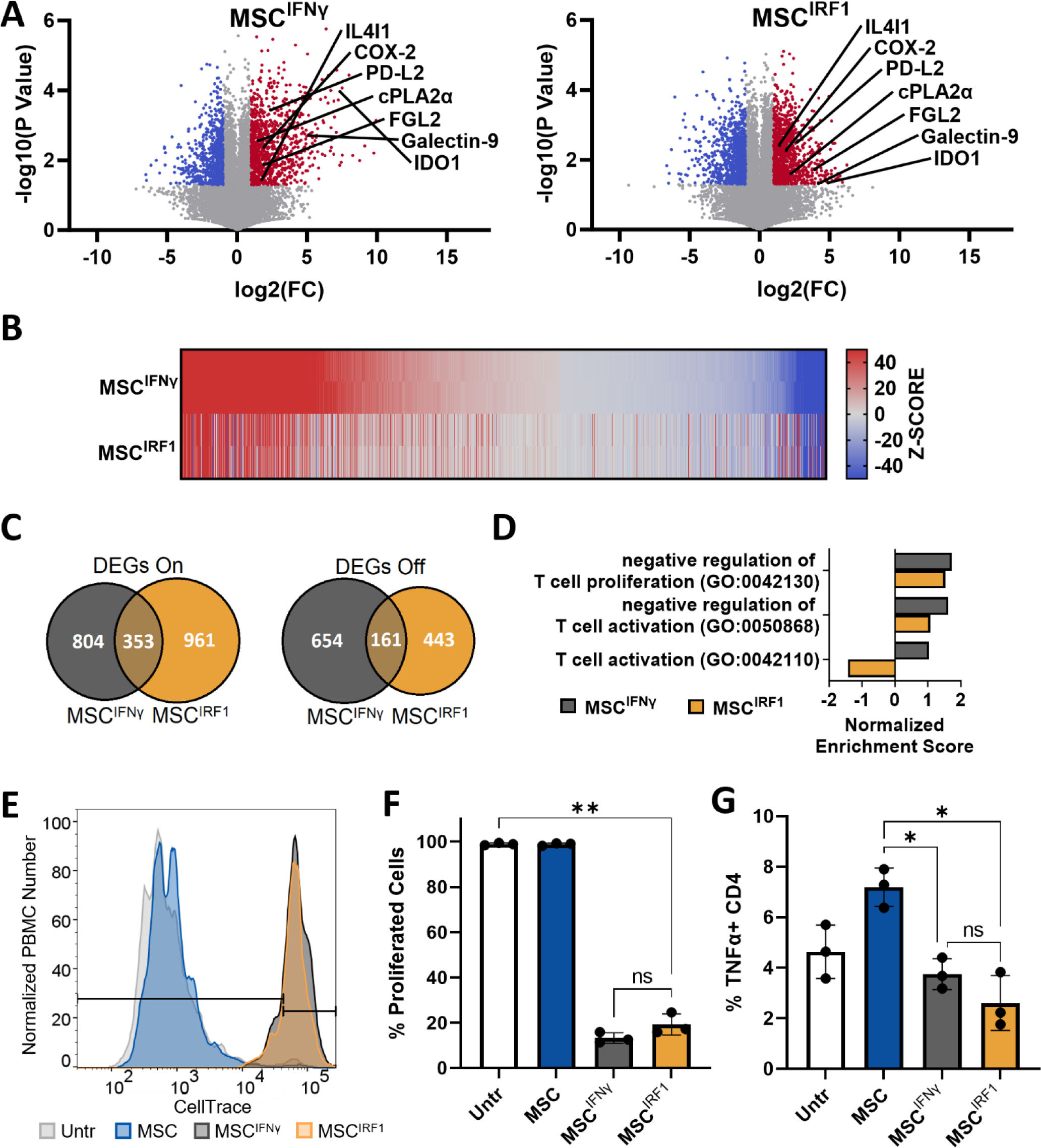
MSC^IRF1^ recapitulates signatures of MSC^IFNγ^ activation and T cell suppression. (A) Volcano plot from RNAseq analyses for MSC^IFNγ^ (left) and MSC^IRF1^ (right). Red and blue regions denote down and up-regulated DEGs respectively (fold change (FC) > 2 and p < 0.05). (B) Expression heatmap for DEGs identified for MSC^IFNγ^. Genes were ordered from highest to lowest z-score. Corresponding z-scores for MSC^IRF1^ are provided. (C) Venn diagram of all DEGs identified for MSC^IFNγ^ and MSC^IRF1^. (D) Normalized enrichment scores for gene ontology (GO) terms associated with T cells suppression / activation. (E and F) Suppression of T cell activation in flow cytometry based, CellTrace Violet (CTV) dilution assays. T cells in PBMC cultures were activated using anti-CD3 and anti-CD28 antibodies and cultured with MSC conditioned media (CM). (G) CD4 T cells were found to express less TNFα on a per-cell basis with CM from MSC^IFNγ^ and MSC^IRF1^ compared to unstimulated MSCs 3 days after exposure to MSC CM. One way ANOVA. *p<0.05, **p < 0.01, ***p < 0.001, ****p < 0.0001 and ns, not significant. Error bars represent standard deviation.

Consistent with their transcriptional profiles, IRF1 and IFNγ-mediated priming were both demonstrated to enhance the immunosuppressive bioactivities of MSCs compared to basal, unprimed cells (Figure 2E-2G, Supplemental Figure S6). MSC conditioned media (CM) prepared using MSC^IRF1^ and MSC^IFNγ^ were shown to suppress the proliferation of peripheral blood monocyte cells (PBMCs) that were pre-stimulated with anti-CD3 and anti-CD28 antibodies to selectively activate T cells (Figures 2E and 2F). Finally, CM from MSCs^IRF1^ was also found to decrease the percentage of CD4 T cells that positively expressed the activation marker TNFα relative to CM from basal, unprimed MSCs (Figure 2G). As with the cell proliferation analyses, similar suppression of CD4 T cells was observed with MSC^IRF1^ and MSC^IFNγ^.

### IRF1 hyperexpression maintains the immune evasive status of MSCs by circumventing STAT1 activation

RNAseq also revealed marked differences between genetic and biochemical priming in the MSC^IRF1^ and MSC^IFNγ^ cells (Figure 2B, 2C, 2D and 3A). In particular, genes from the GO term ‘T cell activation’ (0042110) were depleted with MSC^IRF1^ but positively enriched with MSC^IFNγ^ (Figure 2C). One of the most upregulated MSC^IFNγ^ genes in this GO term was CD74, or HLA Class II Histocompatibility Antigen γ Chain CD74, involved in the assembly and trafficking of MHC class II complexes.^48^ The expression of the transcriptional regulator *CIITA*, which again drives the downstream expression of MHC class II genes, was also significantly upregulated in MSC^IFNγ^ (FC=126±27; p=0.0002) but was much less so in MSC^IRF1^ (FC=6±1.4; p=0.008) (Figure 3A). In turn, while nearly all MHC class II genes were upregulated by IFNγ treatment, HLA gene expression was largely unchanged by IRF1 overexpression (Figure 3A). Consistent with this behavior, flow cytometry analyses of the MHC class II cell surface receptor HLA-DR confirmed this result (Figures 3B and 3C). Here, while the level of HLA-DR was elevated slightly, compared to naïve, unmodified MSCs, the geometric mean of HLA-DR expression in MSC^IFNγ^ was more than 35x that of MSC^IRF1^.

**Figure 3.**
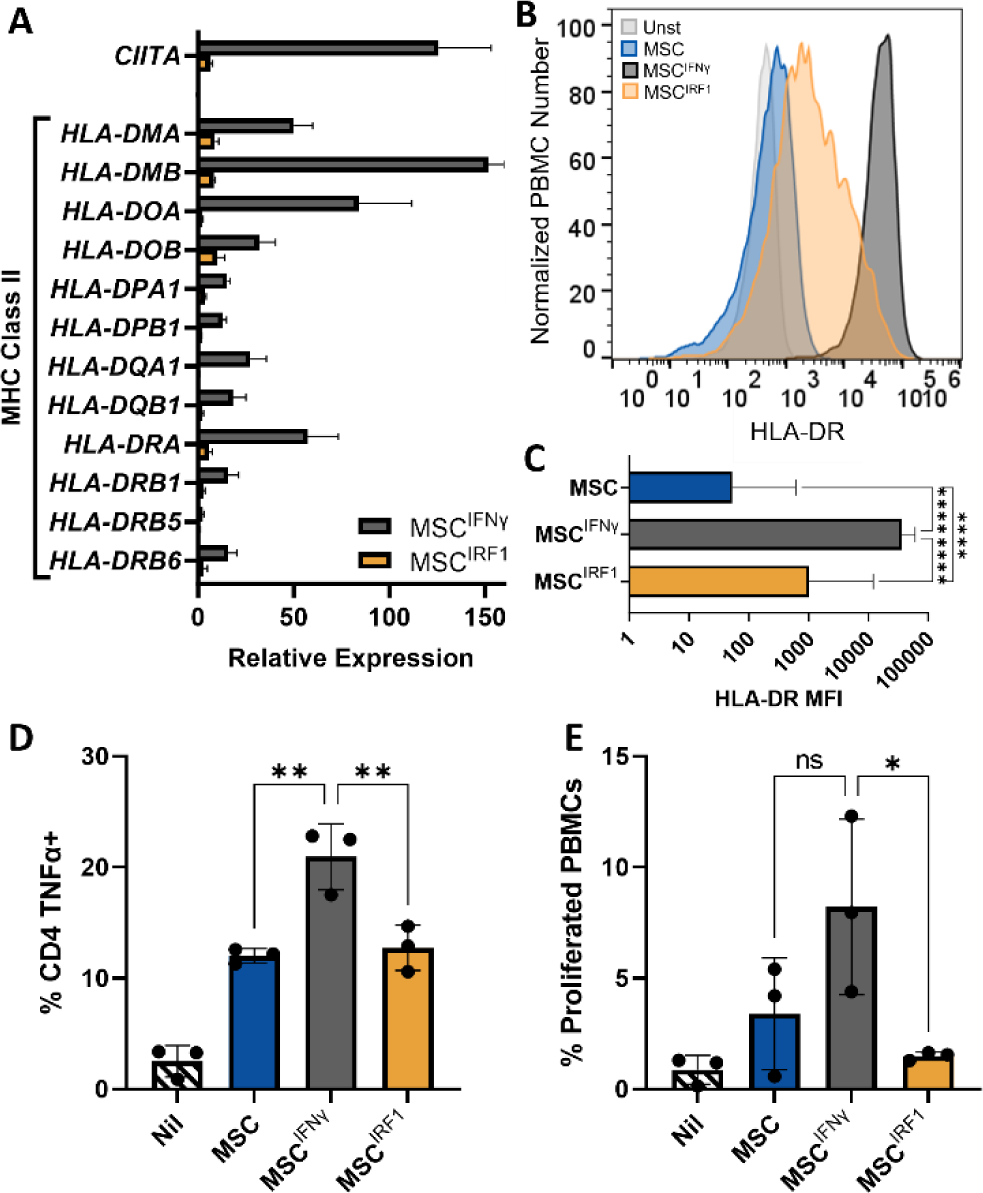
MSC^IRF1^ avoids MHC class II expression and subsequent CD4 T cell activation and proliferation present with MSC^IFNγ^. (A) Relative expression levels of CIITA and MHC class II genes for MSC^IFNγ^ and MSC^IRF1^ with respect to unprimed MSCs. (B and C) Flow cytometry analyses of HLA-DR surface expression. (D and E) Quantitation of TNFα expression in CD4+ T cells (D) and PBMC proliferation (E) upon direct co-culture with MSC^IFNγ^ than MSC^IRF1^ cells. Percent proliferation values were calculated using CellTrace Violet intensities. One way ANOVA. *p<0.05, **p<0.01, ***p<0.001, ****p<0.0001 and ns, not significant. Error bars represent standard deviation.

To test if the above MHC class II expression changes would result in changes in immune reactivity, MSCs, MSCs^IFNγ^, and MSCs^IRF1^ were next co-cultured in direct contact with unstimulated PBMCs. After 7 days, CD4^+^ T cells showed an increase in activation status with MSC^IFNγ^ compared to MSC or MSC^IRF1^ (Figure 3D). There were 60% more TNFα+ CD4 T cells with MSC^IFNγ^ compared to MSC^IRF1^ and 70% more with MSC^IFNγ^ compared to untreated MSC. The PBMCs also proliferated 5.5x more with MSC^IFNγ^ compared to MSC^IRF1^ and 2.4x more with MSC^IFNγ^ compared to untreated MSC (Figure 3E).

### IRF1 priming circumvents STAT1 activation

To gain mechanistic insight into the differences between genetic and biochemical priming, we next examined levels of STAT1 transcription, phosphorylation, and nuclear translocation in naive MSCs, MSC^IRF1^, and MSC^IFNγ^ (Figure 4). IFNγ was found to induce a much larger change in STAT1 transcription (FC=11±0.35, p=0.0005) compared to IRF1 overexpression (FC=3.0±0.39, p=0.02) (Figure 4A). Moreover, IFNγ was found to induce significant STAT1 phosphorylation by western blot as expected (Figure 4B, Supplemental Figure S7).^15^ In contrast, IRF1 overexpression resulted in much less STAT1 phosphorylation. IFNγ also induced much more prominent pSTAT1 nuclear localization in immunocytochemical imaging experiments (Figure 4C and 4D). Together, these results suggest that the IRGs that are differentially expressed in MSC^IRF^^1^ cells relative to naïve MSCs are indeed predominantly activated by IRF1 expression. They further suggest that differences between MSC^IRF1^ and MSC^IFNγ^ transcriptional profiles stem from the ability of IRF1 overexpression to bypass STAT1 phosphorylation and avoid upregulation of STAT1-responsive ISGs that are not activated by IRF1 alone, such as CIITA.

**Figure 4.**
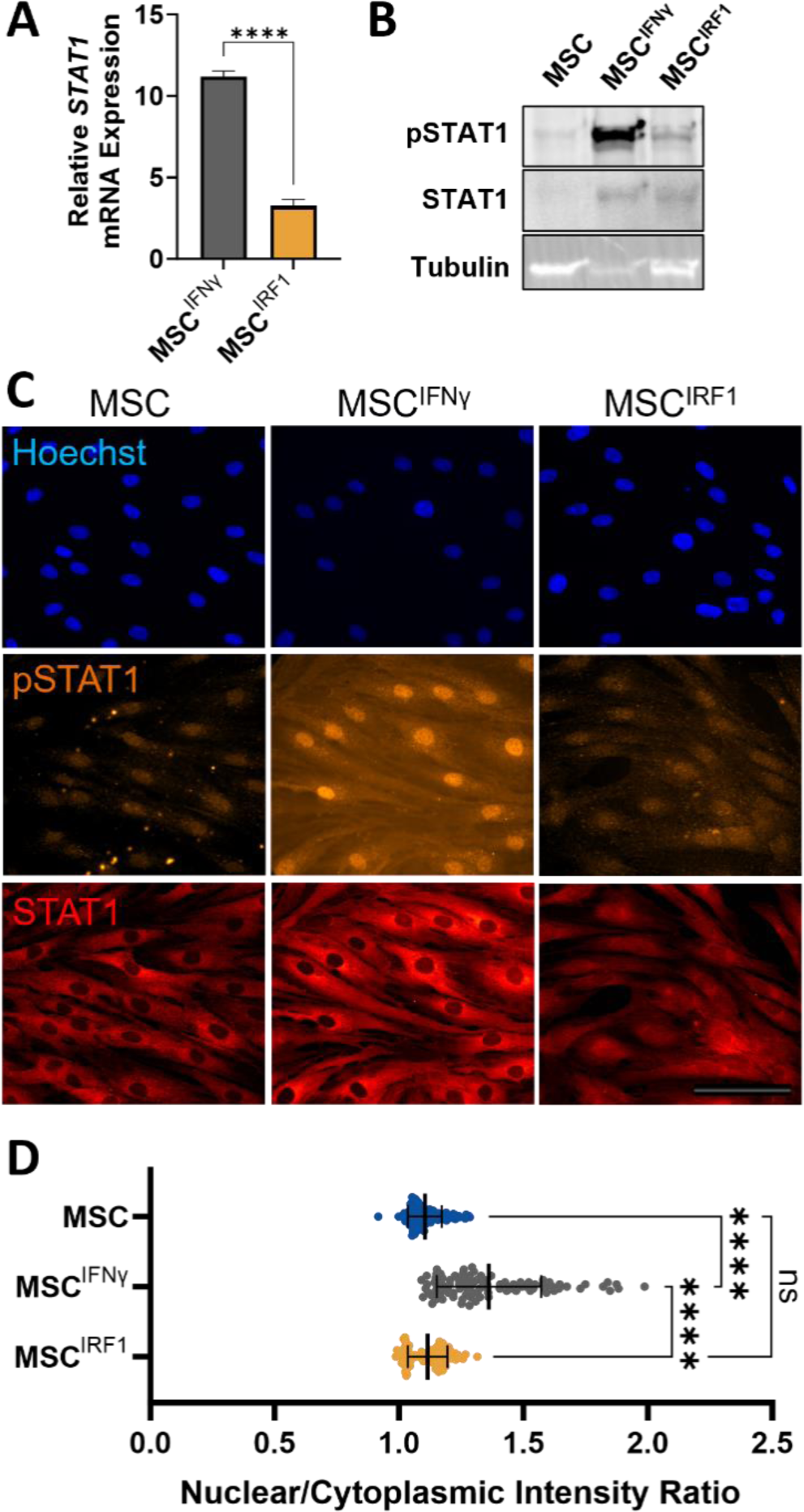
MSC^IRF1^ avoids STAT1 phosphorylation and subsequent signaling present in MSC^IFNγ^ in favor of nuclear localization of unphosphorylated STAT1. (A) Activation of STAT1 transcription by IFNγ stimulation and IRF1 transgene overexpression. (B) Western blotting confirms STAT1 phosphorylation occurs in MSC^IFNγ^, but not MSC^IRF1^ or unprimed MSCs. (C) Immunocytochemical imaging of STAT1 and pSTAT1 localization. Images are contrasted equivalently. Scale bar = 100μm. (D) Summary quantification of nuclear localization of pSTAT1. One way ANOVA. Error bars represent standard deviation.

To further investigate functional consequences of the different activation modes in MSC^IRF1^ and MSC^IFNγ^ cells, high throughput imaging analyses were performed to quantitatively compare differences in primed cell morphologies. Prior reports have shown IFNγ can induce morphological changes that coincide with enhancement in immunosuppressive capacity.^49^ We performed similar analyses using ER tracker dye to stain the MSC endoplasmic reticulum (ER), which extends throughout the cytoplasm, in addition to nuclear staining (Supplementary Figure S8). Seven MSC morphological features were found to be significantly changed upon IFNγ stimulation in this study. Six of these were replicated from previously documented morphological changes in literature, and one of these was not previously studied. Significantly altered features included increases in ER compactness, convex area, perimeter, major axis length, and max feret diameter and decreases in ER form factor and nuclear to ER area ratio. MSC^IRF1^ morphology remained statistically similar to basal MSCs.

### IRF1 can potentiate primary human MSCs activation persistently

We next examined the utility of IRF1 overexpression to sustainably activate the immunosuppressive properties of primary adipose-derived MSCs (AD-MSCs). Here, IRF1 overexpression was again observed to induce IDO1 expression and MSC production of kynurenine (Figure 5A-B) Moreover, conditioned media from IRF1 primed AD-MSCs was also capable of suppressing T cell proliferation in CellTrace dilution assays (Figure 5C-D, Supplemental Figure S9). This suppression was accompanied by reduced activation of CD8+ and CD4+ T cells, signified by a loss of expression of IFNγ and TNFα, respectively (Figure 5E-F). Importantly, the conditioned media in these experiments were also notably prepared using 5-fold fewer cells compared to the hTERT-MSC media, indicating that primary cells are not only amenable to IRF1-medated priming, but higher potencies can be achieved.

**Figure 5.**
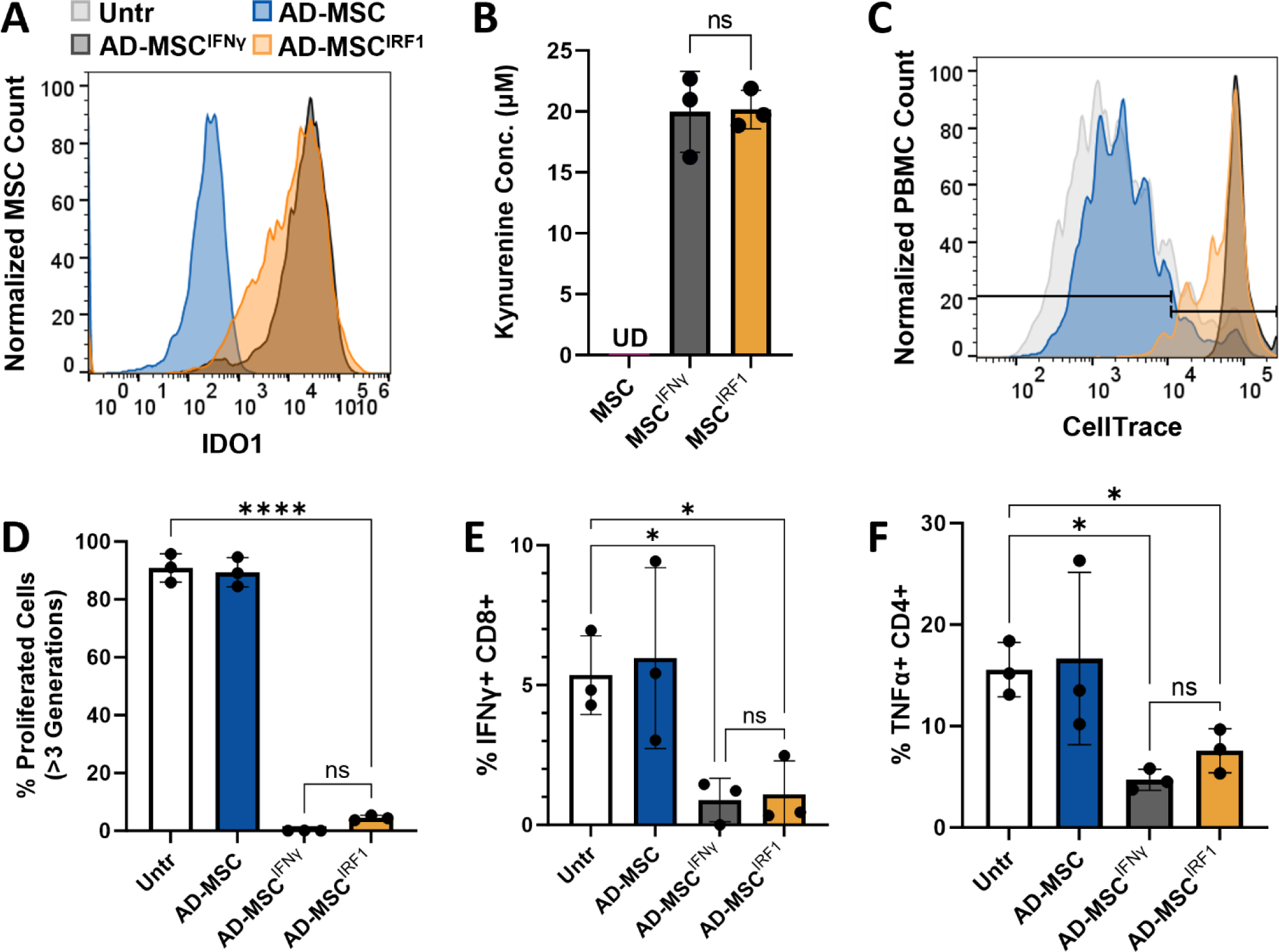
Conditioned media from primary MSC^IFNγ^ and MSC^IRF1^ abrogate T cell proliferation and activation. (A) IDO1 expression measured by flow cytometry shows IRF1 expression results in nearly as much IDO1 expression as IFNγ stimulation. (B) Kynurenine production, an indicator of MSC activity, was high and indistinguishable between MSC^IFNγ^ and MSCIRF1. (C) MSC^IFNγ^ abrogated T cell proliferation slightly more than MSC^IRF1^. Severe proliferation of more than 3 generations was abrogated by both MSC^IFNγ^ and MSC^IRF1^. (D) Summary graph of (panel C). (E) MSC^IFNγ^ and MSC^IRF1^ were similar in their ability to minimize CD8 T cell activation via IFNγ expression. (F) MSC^IFNγ^ and MSC^IRF1^ were similar in their ability to minimize CD4 T cell activation via TNFα expression. One way ANOVA. *p<0.05, **p<0.01, ***p<0.001, ****p<0.0001 and ns, not significant. Error bars represent standard deviation.

A second sample of commercially sourced primary adipose-derived MSCs notably exhibited much lower activation with respect to IDO1 expression (Supplemental Figure S10). IRF1-mediated priming responses were also attenuated appreciably in primary MSCs that were frozen and banked over short timescales (<7 days) (Supplemental Figure S10), indicating IRF1 activation can vary by MSC preparation and patient source. Nevertheless, IDO1 expression remained highly responsive to IFNγ stimulation in these cells. We thus next tested whether IRF1-mediated genetic priming responses could be improved via brief stimulation with IFNγ and/or budesonide, a glucocorticoid that is well known to potentiate MSC responses to IFNγ^50^ (Figure 6A). Our banked primary MSCs were transduced with IRF1 lentivirus, treated with IFNγ and/or budesonide for 24 hours, and then monitored by flow cytometry for IDO1 expression for 21 days. In contrast to biochemical priming alone (Figure 6B), all combinations of IFNγ and/or budesonide with IRF1 overexpression were capable of yielding persistent IDO1 expression for at least 21 days (Figure 6C). Either IFNγ or budesonide was found to increase the rates of IDO1 expression at early time points, indicating potentiation of IRF1 transcriptional activity. Finally, while all treatments that included IFNγ were found to upregulate the expression of HLA-DR, this was not the case with budesonide (Supplemental Figure 11). Instead, the combination of IRF1 priming with budesonide was able to maintain the low levels of HLA-DR expression observed with basal, unprimed MSCs.

**Figure 6.**
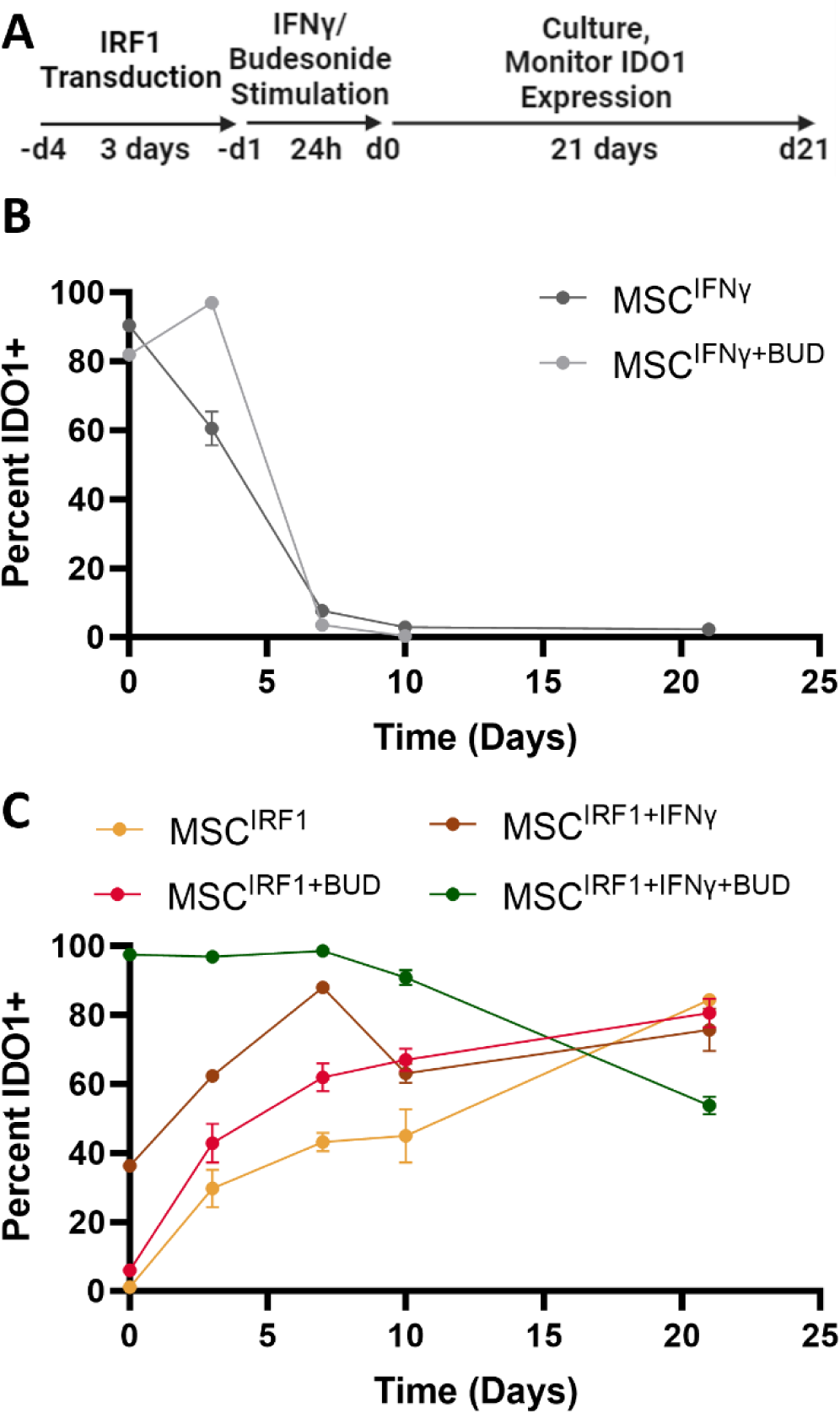
Persistence of IRF1 mediated MSC activation. (A) Experimental scheme for MSC transduction and transient treatment with IFNγ and/or budesonide. (B) IDO1 expression responses in non-engineered MSCs that were primed biochemically with 50ng/mL IFNγ and/or with 1μg/mL budesonide (BUD) for 24 hours (C) The effect of IFNγ and/or budesonide on the activation and persistence of IRF1-transduced MSCs. Percent positive IDO1 cells were determined by flow cytometry. Error bars represent standard deviation, n = 3.

## DISCUSSION

MSCs remain an attractive tool for cell therapy due to their unique abilities to secrete arrays of pleotropic immunomodulatory and restorative biomolecules. MSCs however require some form of biochemical or biophysical priming to activate the production of these molecules since they are not expressed appreciably without stimulation. Priming via pro-inflammatory cytokines such as IFNγ to mimic the reciprocal interactions of MSCs with activated immune cells have been shown to induce broad changes to the MSC transcriptome and secretomes, and improve MSC potency^15,16^. Nevertheless, as demonstrated here and by others, IFNγ-induced potency enhancements are very short-lived and only last a few days.

This work demonstrated constitutive overexpression of the stimulus-responsive transcription factor IRF1 can persistently activate MSC expression of many IFNγ-responsive anti-inflammatory genes, including key immunomodulatory genes like *IDO1* and *PTGS2* (COX-2), as well as other negative regulators of T cell activation. Consistent with this profile, IRF1 overexpression MSCs were capable of suppressing T cell proliferation and activation in both hTERT-modified and primary human MSCs, yielding near equivalent potency found with IFNγ priming.

IRF1-mediated priming was also found to circumvent the expression of the transcriptional activator CIITA, which is upregulated by IFNγ and drives the expression of MHC class II genes known to potentiate MSC immunogenicity and clearance. We attribute this distinction to the role STAT1 plays in activating CIITA transcription, and the utility of IRF1 overexpression to circumvent the phosphorylation of STAT1 in the JAK/STAT pathway. While the relative roles of STAT1 and IRF1 in the upregulation of CIITA are found to vary depending on cell type^51^, the substantial activation of CIITA and downstream MHC class II genes by IFNγ suggests STAT1 dominates this activation in MSCs. The overexpression of IRF1 was not found to influence STAT1 phosphorylation or nuclear localization appreciably via immunocytochemistry or western blotting analyses. Transient IRF1 expression has been reported to promote STAT1 activation in HEK cells^52^. However, our data does not support such effects in MSCs. Instead, the exogenously expressed IRF1 in the genetically primed MSC^IRF1^ cells appears to operate largely independently of the canonical STAT/JAK pathway, enabling selective activation of IRF1-responsive genes without upregulating STAT1.

We also demonstrated IRF1-mediated genetic priming can boost the immunomodulatory bioactivities of primary human MSCs. Here, the potency of conditioned media (CM) from primary MSC^IRF1^ cells proved more potent compared to CM from hTERT modified MSC^IRF1^, with approximately 5-fold fewer cells needed to achieve suppression effects equivalent to those with transient IFNγ stimulation. Nevertheless, we also observed variability in IRF1-mediated priming responses across commercial MSC preparations, donor samples and after freeze-thaw cycles. Responses were measured with respect to IDO1 expression – a key signature of MSC potency. Despite confirmation of efficient lentiviral transduction, IRF1-mediated induction of IDO1 appears to be at times delayed in these cells. In contrast, the onset of IDO1 expression in primary MSCs was notably both rapidly and highly responsive to IFNγ priming in all experiments, despite its subsequent decay.

Building on the hypothesis that the delayed MSC activation response stemmed from potential epigenetic effects that restrict IRF1 access to IRG loci in the absence of complete JAK/STAT signaling, we demonstrated efficient and persistent MSC activation can be achieved by combining IRF1-mediated genetic priming and brief stimulation with IFNγ and / or pharmacological treatment using the corticosteroid budesonide. Both these transient treatments proved capable of accelerating MSC activation (IDO1 expression) appreciably. Moreover, constitutive IFR1 transgene expression was then able to take over and sustain IDO1 activation for at least 21 days; 4-fold longer than IFNγ or IFNγ plus budesonide. Finally, while IFNγ stimulation was naturally found to upregulate HLA-DR expression, the combination of budesonide and IRF1 overexpression minimized this activation, suggesting this combination still circumvents activation of STAT1-responsive ISGs.

MSC treatments with budesonide and other glucocorticoids have been shown previously to enhance IFNγ-induced IDO expression across multiple donors and in over-passaged cells^50^. In these studies, activation improvements were notably maximized with continuous exposure to budesonide. Our study shows that constitutive IRF1 expression can sustain this effect without additional pharmacological treatments. Additional studies are needed to optimize the utility of this approach across different MSC sources and validate the extent to which persistent genetic priming can boost MSCs efficacy *in vivo*. Nevertheless, we anticipate priming via manipulation of IRF1 and/or other stimulus-responsive TFs, individually and in combination, will ultimately facilitate tailored programming of the immunomodulatory and regenerative properties of the secretomes and associated modes of action of MSCs in cell therapy.

## MATERIALS AND METHODS

### MSC culture

Immortalized human adipose hTERT-MSCs and primary adipose MSCs were obtained from ATCC (Manassas, VA). hTERT-MSCs were maintained between passage 2 and 15 in Dulbecco’s modified Eagle’s medium (DMEM; Gibco, Carlsbad, CA) with 10% Fetal Bovine Serum (FBS; Corning, Corning, NY) and 1% antibiotic-antimycotic (Gibco, Carlsbad, CA). Primary MSCs were maintained between passage 2 and 8 in Mesenchymal Stem Cell Basal Medium for Adipose, Umbilical, and Bone Marrow-derived MSCs (PCS-500-030, ATCC, Manassas, VA) with Mesenchymal Stem Cell Growth Kit for Adipose and Umbilical-derived MSCs - Low Serum (PCS-500-040, ATCC) Passaging was done using 0.25% trypsin for 5 minutes at 37°C. Primary MSCs were thawed immediately from the manufacturer and used in downstream experiments unless otherwise specified. In freeze/thaw experiments, primary MSCs were frozen at passage 3 in FBS with 10% dimethyl sulfoxide (DMSO) before being thawed. MSCs were treated with human recombinant IFNγ (PeproTech, Cranbury, NJ) at a concentration of 50ng/mL for 24 hours where indicated.

### Lentiviral MSC engineering

The human IRF1 viral vector was synthesized by VectorBuilder (Chicago, IL). HEK293T cells were obtained from ATCC (Manassas, VA) and maintained in the same media as the hTERT-MSCs between passage 2 and 30. The HEK cells were plated at 7.7E4 cells/cm^2^ the day before transfection. For each transfection, 4.878μg of IRF1 lentiviral plasmid, 1.463μg of pMD2.G and 3.659μg of psPAX2 (Addgene, Watertown, MA) were transfected using JetPRIME (Polyplus, Illkirch, France). Media was exchanged for media containing 4mM sodium butyrate after 8 hours. Virus-containing media was harvested at 24 and 48 hours, filtered with a 0.45μm filter to remove cellular debris and concentrated (100x) using Lenti-X concentrator (Takara, San Jose, CA). The resulting particles were resuspended in DMEM with 10% FBS, 1% antibiotic-antimycotic and stored at −80°C.

The IRF1 lentivirus was reverse transduced into immortalized or primary MSCs in a 6-well plate. MSCs between passage 3 and 5 were added onto the lentivirus and 10mM polybrene was added to each well. Cells were then incubated for 72 hours, their media was changed, and they were incubated for another 3 to 72 hours before their GFP production was assessed by flow cytometry to determine transduction efficiency. A MOI of 1 was the minimum viral volume to obtain approximately 100% GFP positive cells. This viral volume (MOI 1) was used in replicate reverse transduction process to make biological replicates of transgenic IRF1 MSCs that were used in downstream experiments. Transduction efficiencies were confirmed via fluorescence imaging.

### RT-qPCR

Immortalized human adipose MSCs were plated in 6 well plates and rested for 24 hours, after which, cells were lysed, and RNA was extracted using a RNeasy Mini Kit (Qiagen, Germantown, MD). RNA was reverse transcribed to cDNA using an iScript cDNA Synthesis Kit (Bio-Rad, Hercules, CA). qPCR was conducted using SsoAdvanced Universal SYBR Green Supermix (Bio-Rad, Hercules, CA) and the following primers (IDT, Newark, NJ):

IRF1 F: 5’-CTCTCCCCGACTGGCACATC-3’

IRF1 R: 5’-CCGACTGCTCCAAGAGCTTCA-3’

IDO1 F: 5’-ACGGGACACTTTGCTAAAGGC-3’

IDO1 R: 5’-GGTTGCCTTTCCAGCCAGACA-3’

GAPDH F: 5’-CAATGACCCCTTCATTGACC-3’

GAPDH R: 5’-TTGATTTTGGAGGGATCTCG-3’

### RNA Sequencing

RNAseq experiments were performed in duplicate. RNA was extracted using an RNEasy Mini Kit and frozen at −20°C and sent to Azenta (Burlington, MA) for whole RNA sequencing. FASTQ files were processed using Galaxy to quantify read counts for each gene and condition. R and Python were used to convert Ensemble gene IDs to gene symbols. Python and the GSEAPY package were used to conduct gene ontology enrichment analyses.

### Western blotting

MSCs were lysed in RNA immunoprecipitation (RIPA) buffer. Total protein concentrations were quantified using a BCA assay. 25μg of protein was loaded for SDS-PAGE and transferred onto a PVDF membrane for western blot. Primary antibodies for STAT1 (66545-1, Proteintech, Rosemont, IL) and pSTAT1 (MA5-15071, Thermo, Waltham, MA) were used at a 1:5000 and 1:1000 dilution, respectively, in 1X Tris Buffered Saline with 1% Casein (Bio-Rad, Hercules, CA). Secondary α-mouse HRP (A6154, Sigma-Aldrich, St. Louis, MO) or α-rabbit HRP (ab6721, Abcam, Cambridge, UK) were used at a 1:3000 dilution in 1X Tris Buffered Saline with 1% Casein. Membranes were exposed after addition of ECL (170-5060, Bio-Rad, Hercules, CA). Tubulin was detected with hFAB™ Rhodamine Anti-Tubulin Primary Antibody (12004166, Bio-Rad, Hercules, CA) at 1:3000 dilution.

### Immunocytochemistry

MSCs were plated in 6 well plates containing coverslips, cultured for 24 hours, and fixed with 4% PFA in PBS for 20min at 4-8°C. Cells were washed thoroughly with 0.2μm filtered 5% BSA in PBS (blocking solution), and permeabilized by incubation in 0.2% Triton-X-100 in PBS at room temperature for 5min. After washing and incubating with blocking solution for 1 hour, cells were immuno-stained overnight at 4-8°C with rabbit anti-human IRF1 antibody (1:1000, ab243895, Abcam, Cambridge, UK) or rabbit anti-human pSTAT1 (1:400, MA5-15071, Thermo, Waltham, MA) and mouse anti-human STAT1 (1:800, 66545-1, Proteintech, Rosemont, IL) diluted in blocking buffer. Secondary antibody staining was performed using Cy5 Goat anti-rabbit (1:500, A10523, Thermo, Waltham, MA), AF 594 donkey anti-rabbit (1:500, A21207, Invitrogen, Waltham, MA), or AF 647 goat anti-mouse (1:500, A21235, LTC, Carlsbad, CA). Secondary staining solutions also contained Hoechst nuclear stain (1:10,000, 33342, Thermo, Waltham, MA). Cells were visualized with a Nikon Eclipse Ti2 at 60x magnification. Background subtraction was done using ImageJ and was performed identically for each fluorophore.

### Flow cytometry

Cells were trypsinized, fixed using BD Cytoperm/Cytofix (BD, Franklin Lakes, NJ), permeabilized with BD Perm/Wash Buffer, and stained at room temperature for 1 hour using a PE labeled mouse anti-IDO1 antibody (12-9477-42, Thermo, Waltham, MA) or a BV786 labeled mouse anti-human HLA-DR antibody (564041, BD, Franklin Lakes, NJ) according to manufacturer instructions. Excess antibody was removed via centrifugation and supernatant removal. Fluorescence levels were then quantified.

### Kynurenine production

Kynurenine concentration was assessed by Ehrlich reaction. 100μL conditioned media was combined with 50μL 30wt% trichloroacetic acid (TCA) in water in a round-bottom 96 well plate. A standard curve of purified L-kynurenine (Sigma, St. Louis, MO) dissolved in R10 media was also included. This mixture was spun down at 2200RCF. 100μL supernatant was combined with fresh 2 wt% dimethylaminobenzaldehyde (Sigma, St. Louis, MO) in glacial acetic acid (Fisher, Waltham, MA). This mixture was immediately read at 490nm on a plate reader.

### Suppression bioactivity assays

MSC conditioned media was prepared by culturing primary MSCs (2×10^5^ cells) or hTERT-MSCs (1×10^6^ cells) in R10 media for 48-72 hours (RPMI Medium 1640 (Gibco, Waltham, MA) with 10% heat-inactivated Fetal Bovine Serum, 1% Penicillin-Streptomycin (Gibco), 1% GlutaMAX Supplement (Gibco), 1% Sodium Pyruvate 100mM (Gibco), 1% HEPES 1M (Gibco), 1% MEM Non-Essential Amino Acids (Gibco) for 48 and 72 hours, respectively. Conditioned media was used directly without storage in subsequent assays. Human PBMCs were prepared from buffy coats obtained from Gulf Coast Regional Blood Center (Houston, TX, USA). Buffy coats were frozen and stored in liquid nitrogen before being thawed just prior to use. Viability was measured by AOPI (Nexcelom, Lawrence, MA) and was required to be >90% at thaw.

T cells were activated by incubating PBMCs with anti-CD3 (OKT3, Tonbo/Cytek, San Diego, CA) and anti-CD28 (302934, Biolegend, San Diego, CA) antibodies in PBS at 37°C for 3 to 5 hours. For T cell proliferation analyses, PBMCs were stained with CellTrace Violet (CTV) (Thermo, Waltham, MA) prior to activation.

TNFα and IFNγ expression levels were measured by treating PBMCs with GolgiPlug (Fisher, Waltham, MA) and GolgiStop (Fisher) to facilitate immunostaining analyses. PBMCs were fixed and permeabilized for 1 hour at room temperature with Foxp3/Transcription Factor Fix/Perm Concentrate and Diluent (Tonbo/Cytek), and then washed with Tonbo Flow Cytometry Perm Buffer (Tonbo/Cytek). Cells were then immuno-stained 1 hour at room temperature with a panel of antibodies: anti-CD3 (563725, BD, Franklin Lakes, NJ), anti-TNFα (Biolegend 502938), anti-IFNγ (Biolegend 506516), anti-perforin (Biolegend 506516), anti-T-bet (Biolegend 644812), and anti-Ki67 (BD 564071) for. Viable cells were identified using Ghost Dye Violet 510 (Tonbo/Cytek). Flow cytometry was performed using a BD LSRFortessa instrument. T cell proliferation and activation were quantified identically in MSC-PBMC co-culture experiments. In these experiments, MSCs were cultured for 3 days before PBMCs were placed in coculture for an additional 7 days. Media was changed every other day.

## DATA AVAILABILITY

All original data is available from the authors under request.

## SUPPLEMENTAL INFORMATION

Supplemental figures are available.

## ACKNOWLEDGEMENTS

We thank the Cytometry and Cell Sorting Core of Baylor College of Medicine for the use of the LSRFortessa Flow Cytometer. We thank the Shared Equipment Authority of Rice University for the use of the Sony MA900 Cell Sorter. We thank BioRender.com for facilitating the creation of the schematics in this manuscript. This work was supported by grants from the NSF (2041107), and NIH (R21EB030772).

## AUTHOR CONTRIBUTIONS

M.A.B. and M.R.D conceived of the project. M.A.B., G.C.B., O.V., P.L.W., A.R.D., I.B.H., and M.R.D. designed experiments. M.A.B., G.C.B., D.A.B., L.D.S.S., T.N., A., S.C.L., and B.M. conducted experiments. M.A.B., G.C.B, L.D.S.S., and M.R.D. analyzed data. M.A.B. and M.R.D. wrote the manuscript. M.A.B., O.V., P.L.W., A.R.D., I.B.H., and M.R.D. edited the manuscript.

