## Supplemental Figures for "Persistent tailoring of MSC activation through genetic priming"

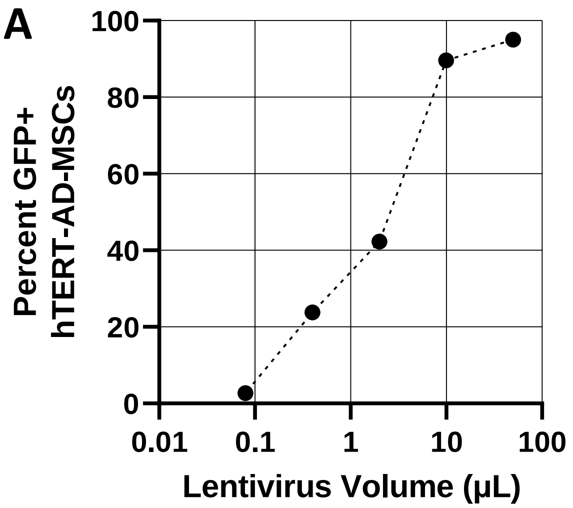


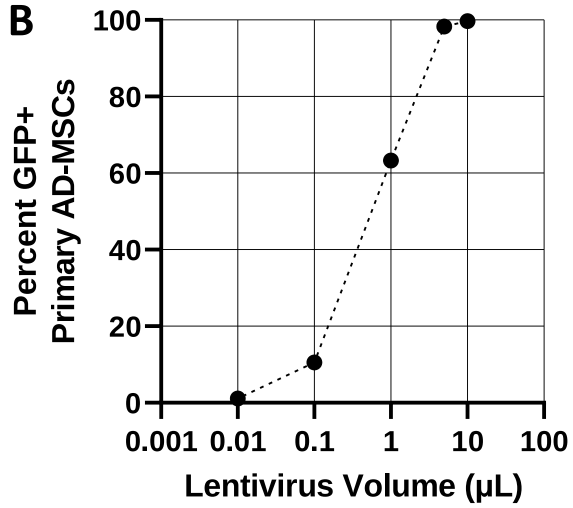


**Figure S1 IRF1 lentivirus titering**

**(A)** Flow cytometry analysis of immortalized human adipose MSCs transduced with varied amounts of hIRF1 lentivirus. **(B)** Flow cytometry analysis of primary human adipose MSCs transduced with varied amounts of hIRF1 lentivirus.

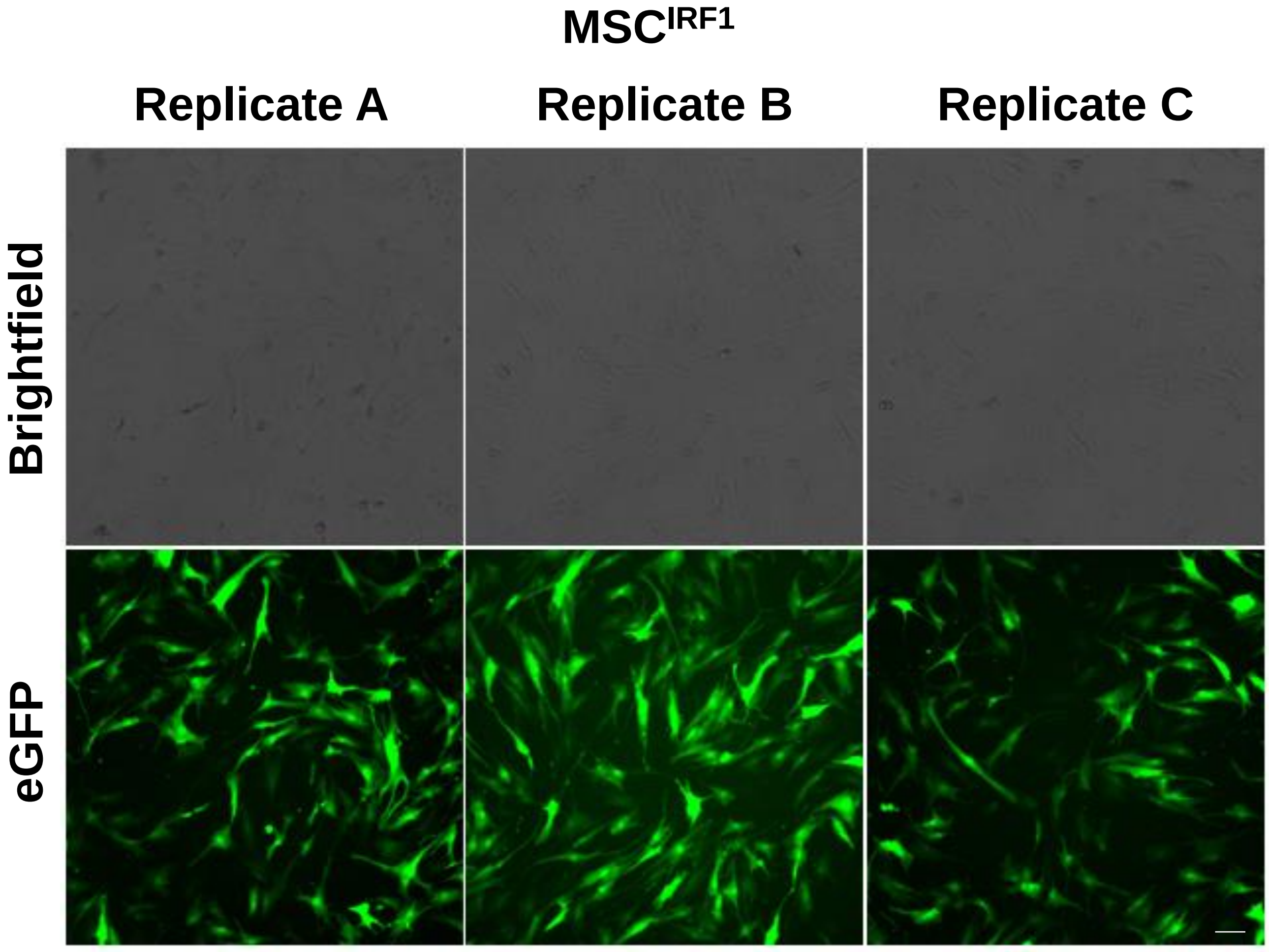


**Figure S2 Fluorescence imaging confirmed immortalized MSC replicates were similar and had an approximate transduction efficiency of 100%**
Lentivirus contained both hIRF1 and eGFP. Replicates pictured were carried forward to all downstream experiments involving immortalized MSCs. Scale bar 100µm.

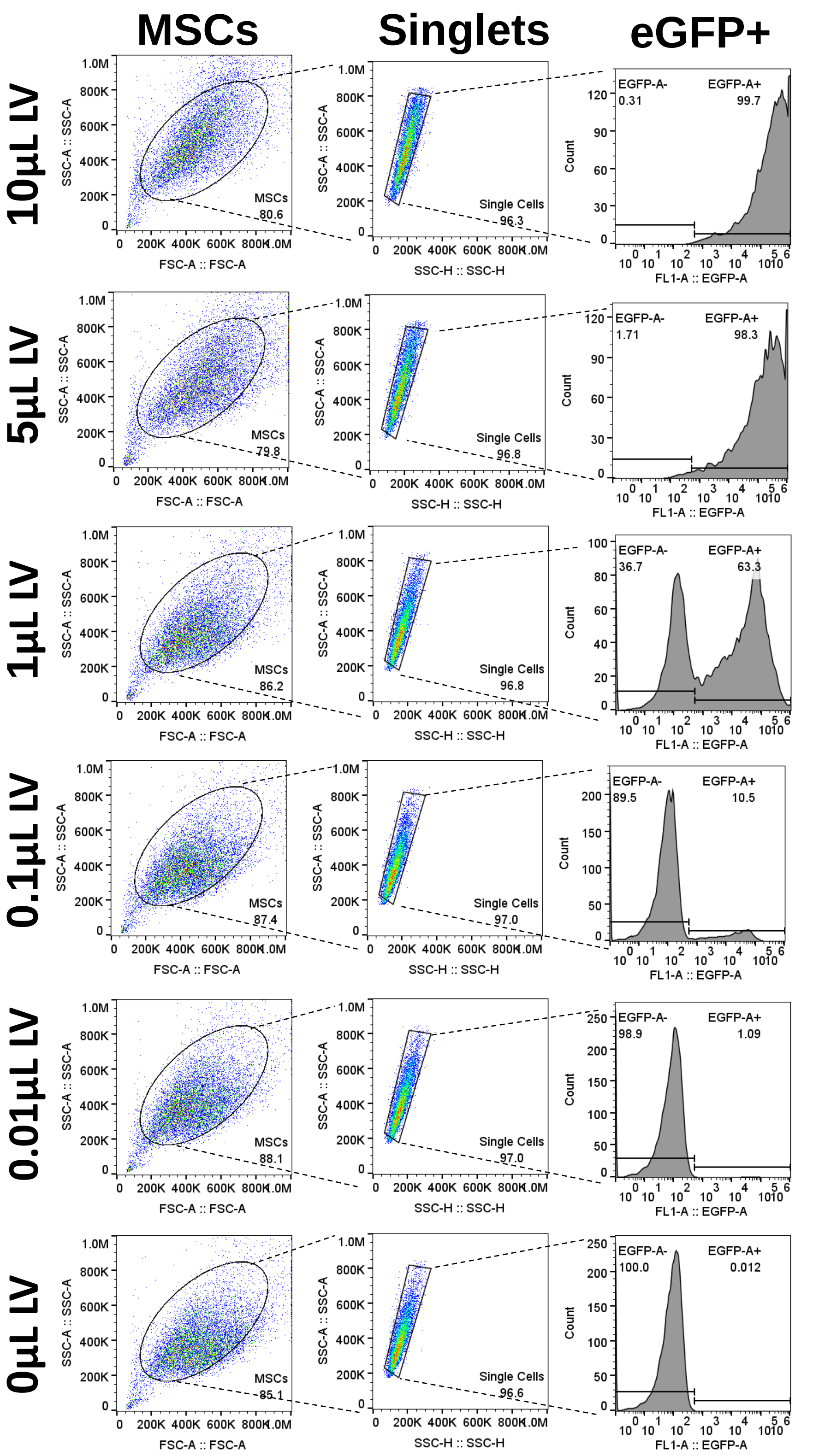


**Figure S3 Exemplar gating and results of IRF1 lentiviral titering**

Flow cytometry analysis of primary human adipose MSCs transduced with varied amounts of hIRF1 lentivirus. Lentivirus contained both hIRF1 and eGFP under separate promoters.

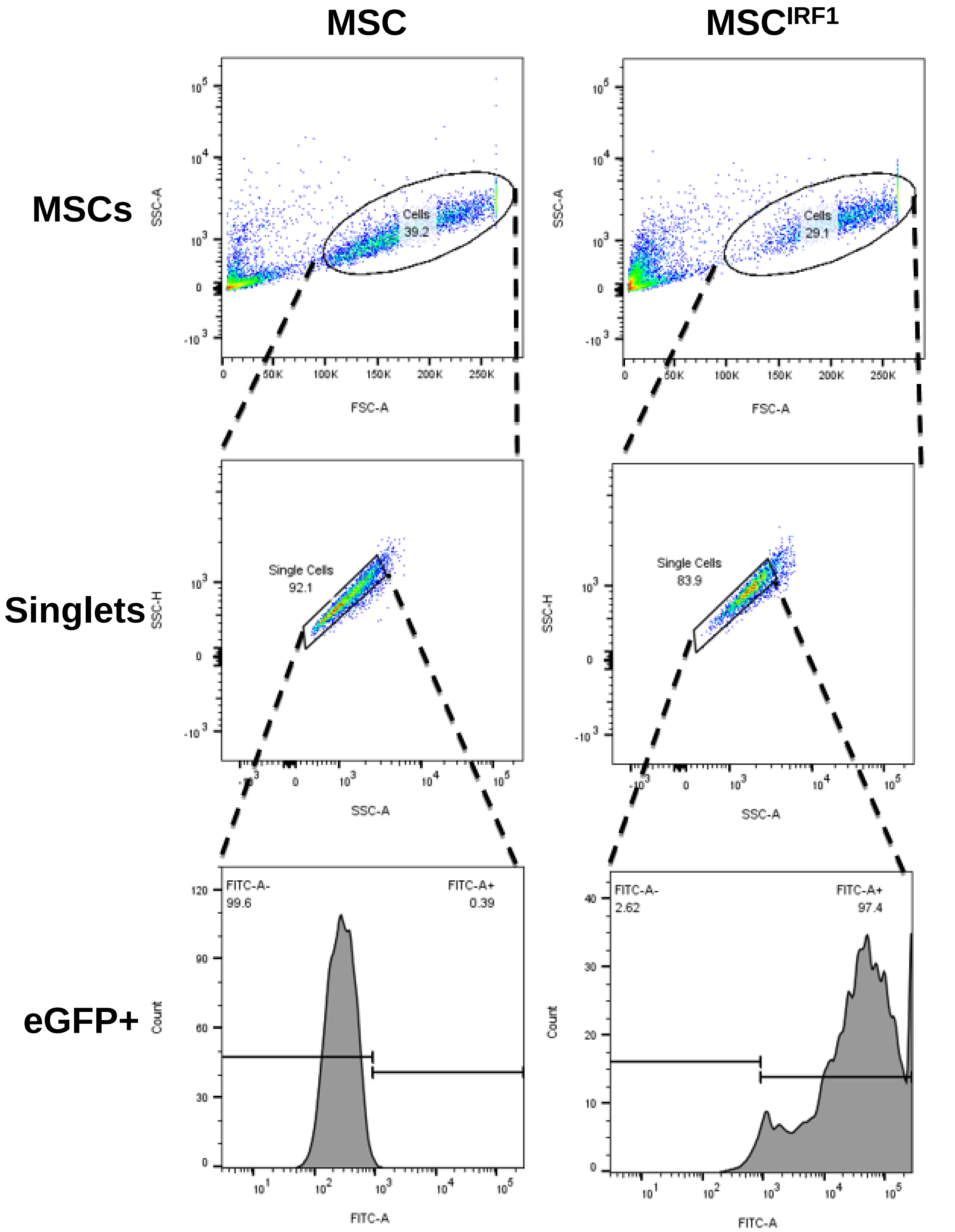


**Figure S4 Exemplar hIRF1 lentivirus transduction of immortalized human adipose MSCs**
Flow cytometry analysis of immortalized human adipose MSCs transduced with IRF1 lentivirus, one of three transduced replicates that were moved forward into all downstream experiments. Lentivirus contained both hIRF1 and eGFP under separate promoters.

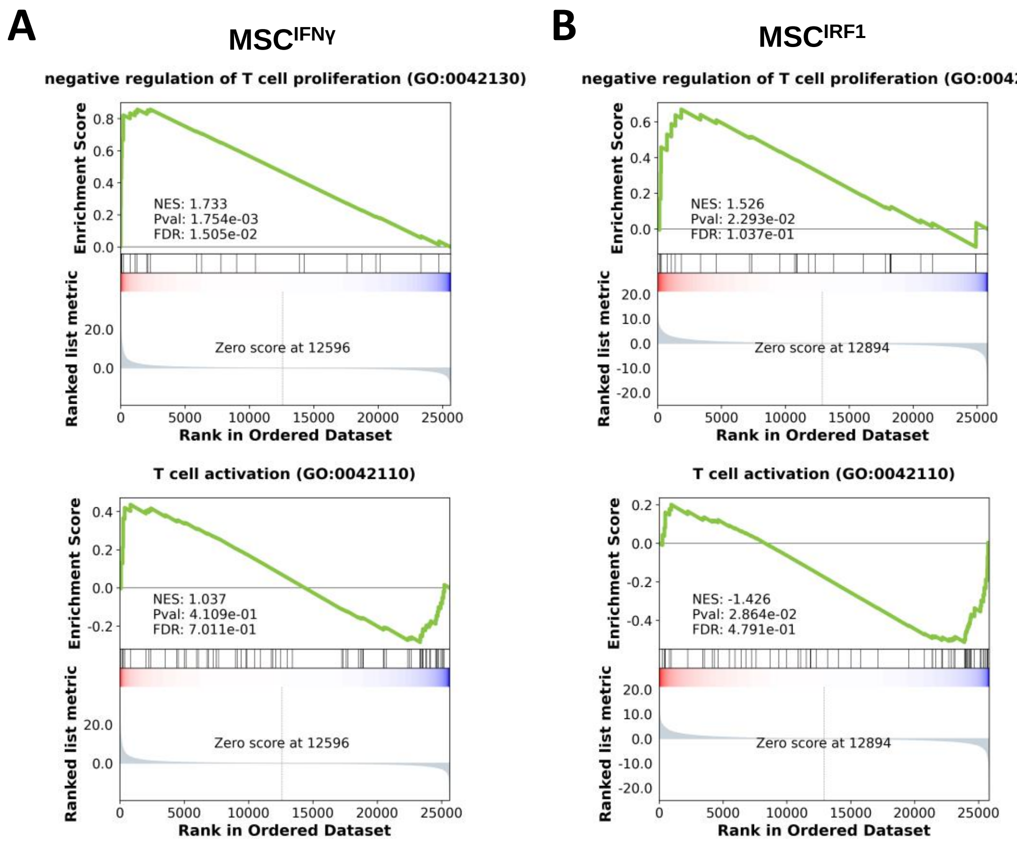


**Figure S5 MSC$^{IRF1}$ avoids T cell activation while largely maintaining ability to negatively regulate T cell proliferation**

GSEA analysis of RNAseq for **(A)** MSC$^{IFN\gamma}$ and **(B)** MSC$^{IRF1}$ for GO terms key to MSC therapeutic function experimentally tested in this work.

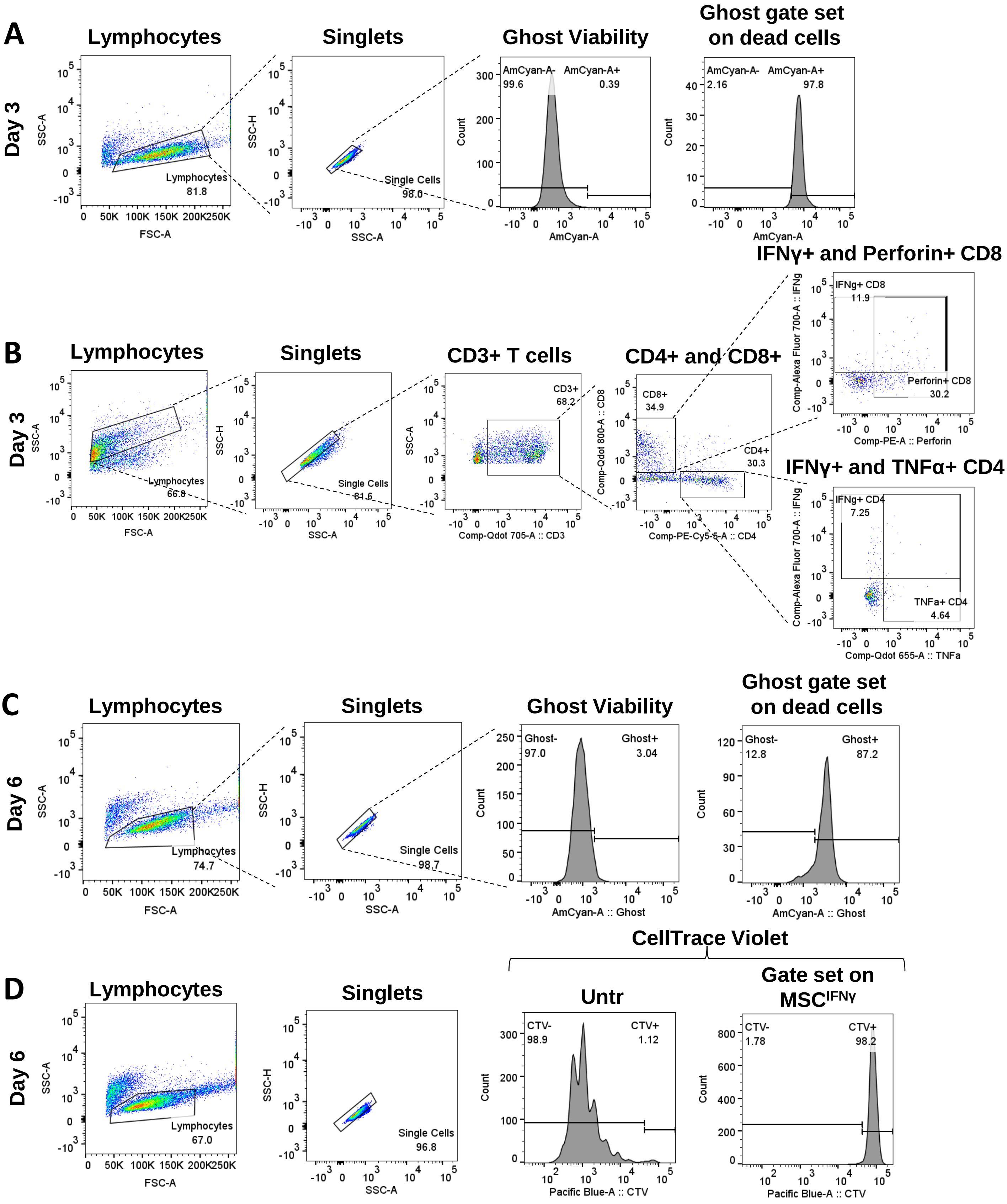


**Figure S6 Flow cytometry gating schemes for PBMCs cultured with conditioned media from immortalized MSCs**

**(A)** Representative day 3 viability of PBMCs via Ghost Violet 510. **(B)** Representative day 3 immunophenotyping. **(C)** Representative day 6 viability of PBMCs via Ghost Violet 510. **(D)** Representative day 6 CTV quantification. Untr = untreated or no MSCs.

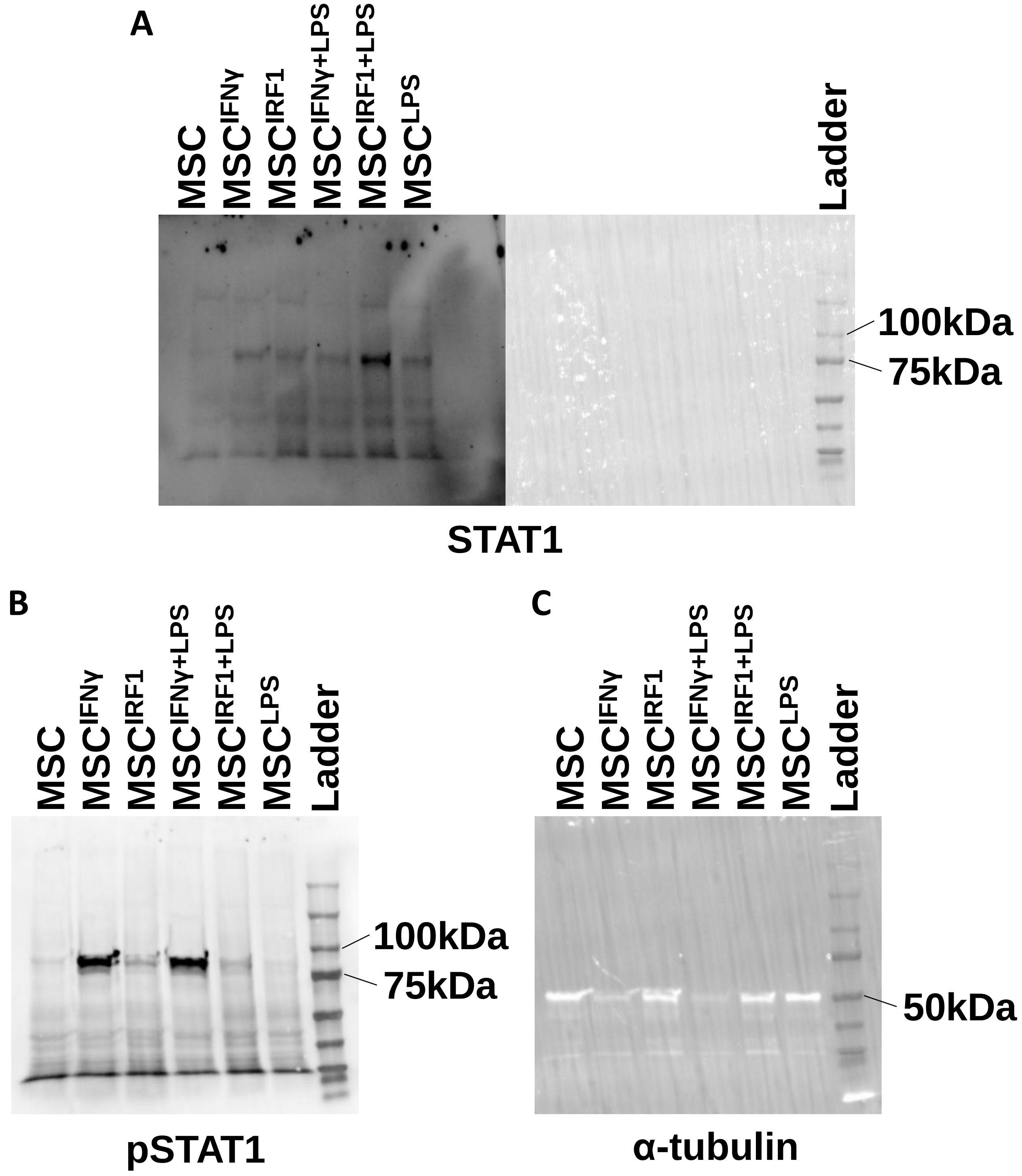


**Figure S7 Original images of western blots**

**(A)** Western blot analysis of STAT1 depicting the same blot with HRP (left) and colorimetric (right) imaging done separately to view the target protein and the protein ladder. **(B)** Western blot analysis of phospho-STAT1 or pSTAT1. **(C)** Western blot analysis of α-tubulin control.

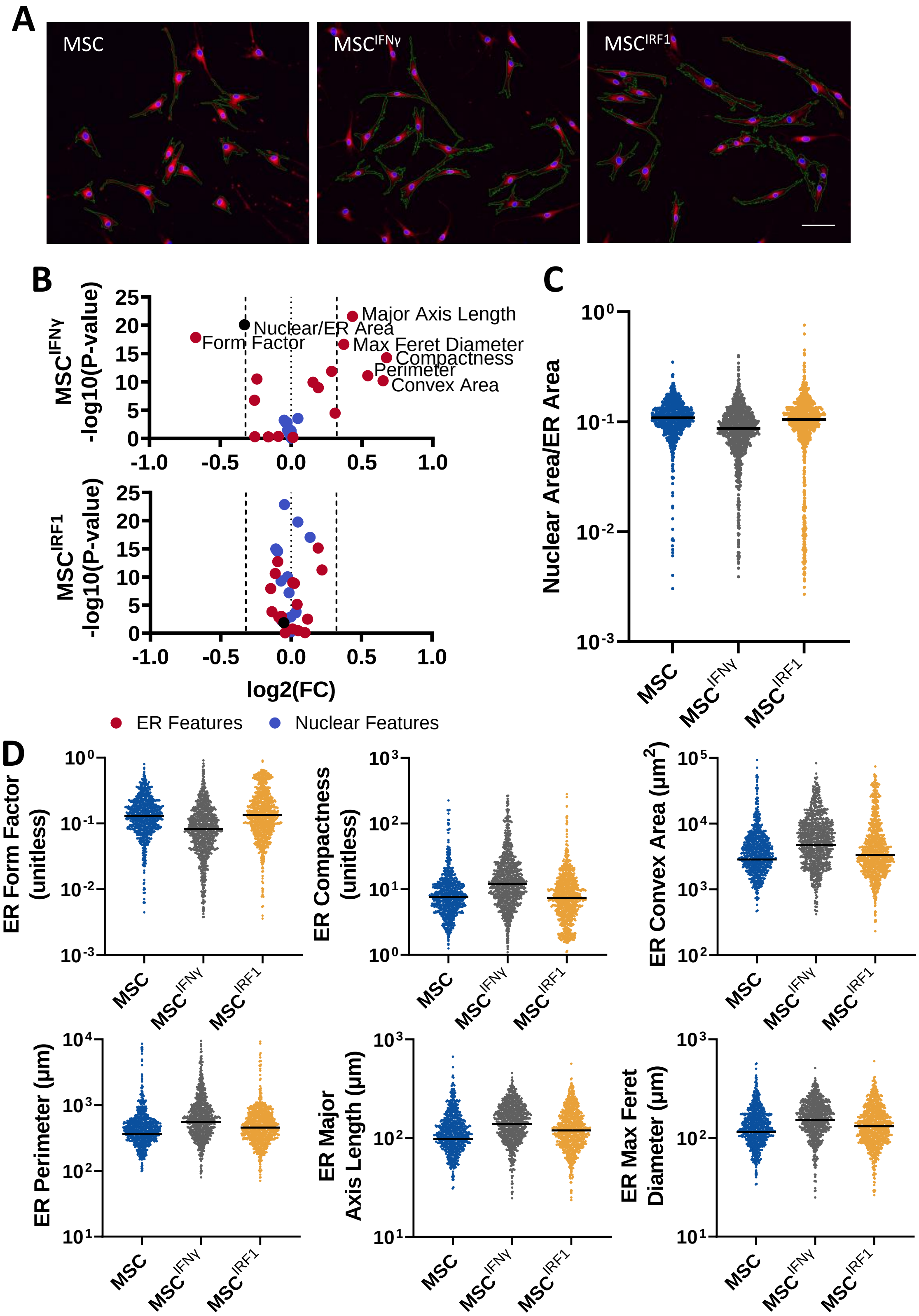


**Figure S8 MSC$^{IRF1}$ does not undergo morphological changes experienced by MSC$^{IFN\gamma}$**

**(A)** Volcano plots of measured morphological features of MSC endoplasmic reticulum (ER), nuclei, or a ratio of the two, both vs MSC, shows MSC$^{IFN\gamma}$ undergoes morphological changes while MSC$^{IRF1}$ does not. Dashed lines set to fold change of 1.25. **(B and C)** Individual cells plotted for the most significantly changing morphological features nuclear to ER area ratio (panel B) and ER features (panel C). **(D)** Exemplar images with red depicting the ER (ER-Tracker Red), blue depicting nuclei (Hoechst), and green indicating the edges of the ER determined using CellProfiler. Scale bar 100μm.

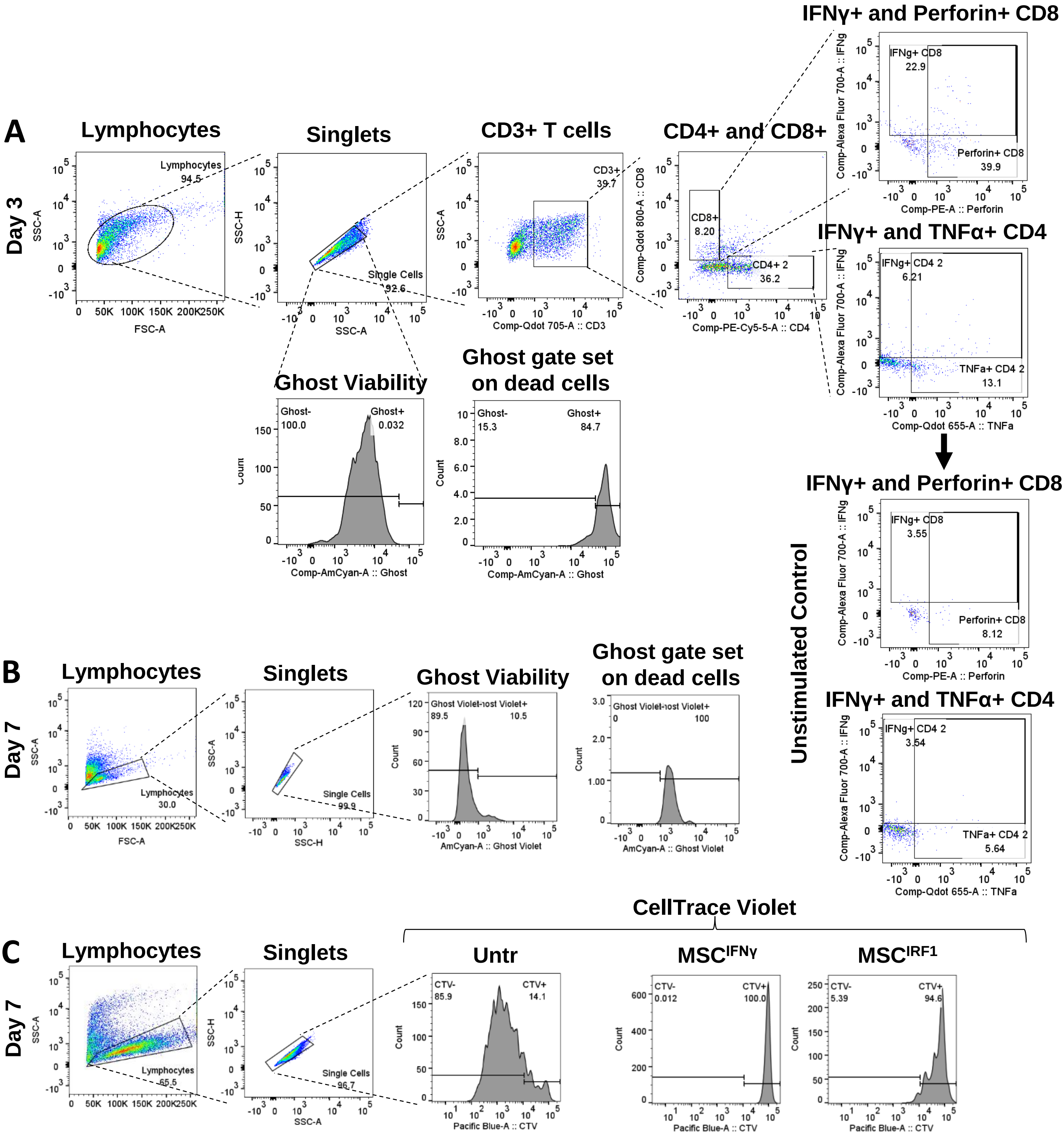


**Figure S9 Flow cytometry gating schemes for PBMCs cultured with conditioned media from primary MSCs**

**(A)** Representative day 3 immunophenotyping, including viability. **(B)** Representative day 7 viability of PBMCs via Ghost Violet 510. **(C)** Representative day 7 CTV quantification. Untr = untreated or no MSCs.

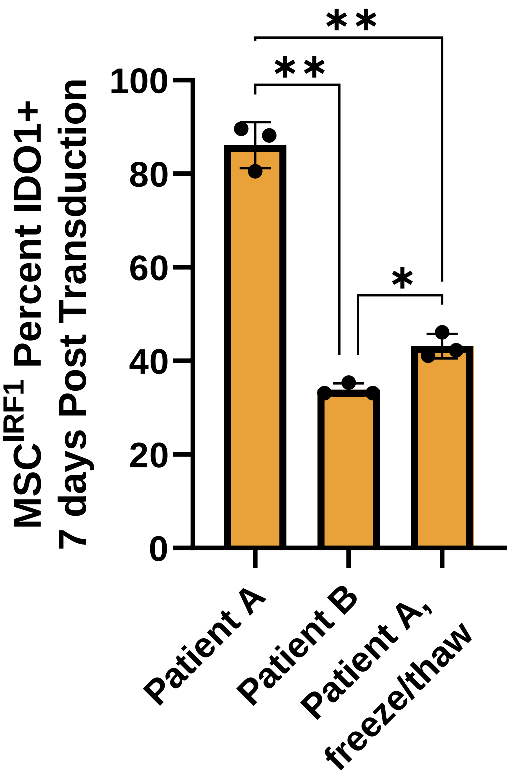


**Figure S10 Patient variability and freeze/thaw cycles negatively affect MSCs' ability to express IDO1 in response to IRF1**
Primary MSCs 7 days following IRF1 transduction, with and without an extra freeze/thaw cycle or with MSCs from two patients

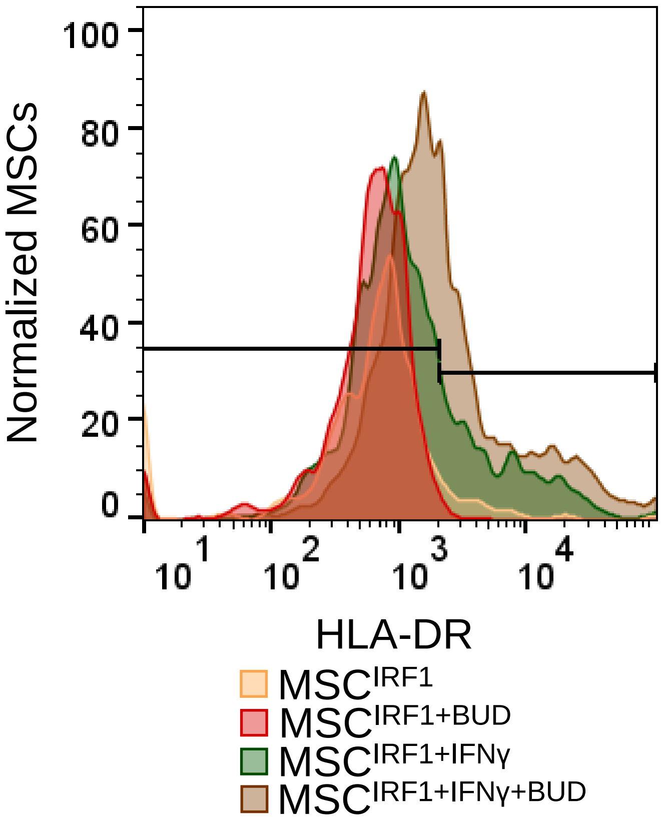


**Figure S11 MHC Class II molecule HLA-DR expression following stimulation to hasten IDO1 activation**
**(A)** Day 10 following IRF1 transduction and 24 hours of stimulation with or without IFNγ, budesonide (BUD) or both.
